# A novel innate pathogen sensing strategy involving ubiquitination of bacterial surface proteins

**DOI:** 10.1101/2021.10.20.465158

**Authors:** Shruti Apte, Smita Bhutda, Sourav Ghosh, Kuldeep Sharma, Osheen Sahay, Jyotirmoy Rakshit, Akash Raj Sinha, Soham Dibyachintan, Suvapriya Roy, Akshay Datey, Shweta Santra, Jincy Joseph, Sreeja Sasidharan, Sven Hammerschmidt, Dipshikha Chakravortty, Manas Santra, Anirban Banerjee

**Author notes:** These authors contributed equally to this work (Shruti Apte, Smita Bhutda, Sourav Ghosh).

## Abstract

Sensing of pathogens by ubiquitination is critical for maintaining cytosolic sanctity. However, universal ubiquitination targets on bacteria, especially of proteinaceous origin, remain unidentified. Here, we unveil a novel strategy, involving recognition of degron-like motifs for identification of first protein-based ubiquitination substrates on phylogenetically distinct bacteria. Such motifs can form a new class of intra-cytosolic pathogen associated molecular patterns (PAMPs) as their incorporation enables identification of non-ubiquitin targets by host Ub-ligases. We find SCF^FBW7^ E3-ligase, supported by the regulatory kinase, GSK3β, is crucial for effective pathogen detection and clearance. This may explain the enhanced risk of infections in Chronic Lymphocytic Leukaemia patients bearing FBXW7 mutations. We conclude that exploitation of such ubiquitous pathogen sensing strategy allows conservation of cellular resources and boost anti-microbial immunity.

**One Sentence Summary:** Ubiquitination of bacterial surface proteins fosters sensing and clearance of diverse pathogens

## Main Text

Surveillance of intracellular milieu for restriction of pathogen proliferation is critical to preserve cytosolic sterility. A breach of such defense mechanisms not only provides the pathogen a refuge from extracellular innate immunity but also offers opportunity for rapid multiplication and dissemination within the host (*1*). Thus, a consortium of potent pathogen sensing mechanisms and cell autonomous defense systems are critical to restrict invasive pathogens. Ubiquitination is one of such ubiquitous strategy which plays a pivotal role in pathogen recognition and elimination (*2*). However, the common tactics adopted by the host for identification of ubiquitination substrate for the variety of pathogens it encounters remains elusive. Since ubiquitination is also critical for maintenance of cellular homeostasis, a universal approach for substrate identification is necessary to create a frugal system for efficient and optimal resource utilization. Recent studies have depicted few secreted effector proteins and lipopolysaccharide as ubiquitination substrates; however, neither their presence is ubiquitous nor they have pronounced role in direct bacterial elimination (*3-5*). In this study, we uncover a simple yet generic principle that the host adopts for sensing phylogenetically diverse pathogens of both Gram-positive as well as Gram-negative origin. Employing this, we not only identified the first bacterial surface protein as a substrate for cellular ubiquitination, further it remarkably facilitated conversion of a non-ubiquitinable surface protein to ubiquitin substrate for efficient bacterial clearance. Such a conserved mechanism of cytosolic pathogen recognition is central to fend off bacterial infections.

Upon sensing cytosolic invasion by pathogens, the host marks them with poly-Ub chains to trigger their clearance. Since such poly-Ub chains are primarily composed of K48 and K63-Ub topologies (*6*), we first explored the predominance and spatial location of these chain types on two phylogenetically distinct pathogens, *Streptococcus pneumoniae* (SPN) and *Salmonella enterica* serovar Typhimurium (STm) which causes pneumonia and gastroenteritis in humans, respectively. For both these pathogens, existence of cytosolic life has been previously described (*7, 8*). Using ubiquitin linkage specific antibodies, we observed that significantly higher intracellular bacteria were marked with K48-Ub chain type (∼26% for SPN and 37% for STm) in contrast to K63-Ub (Fig.1A). Analysis of spatial location by Structured Illumination Microscopy (SIM) indicated that cytosolic (free of vacuolar remnants) or cytosol exposed bacteria (within damaged endosome) are primarily associated with K48-Ub while K63-Ub signal was located on damaged endosome which were marked with Galectin-8 (Gal8, endosome damage sensing marker) (*9*) (Fig.1B-E). Indeed, ∼99% and ∼76% of K48-ubiquitinated SPN and STm, respectively, were devoid of Gal8 (Fig.1F). Collectively, these suggested that coating of bacterial surface with K48-Ub chains is a major pathogen sensing mechanism employed by the host for recognizing cytosol dwelling bacteria.

We next attempted to identify the substrate for K48-ubiquitination on the bacterial surface. As host E3 ubiquitin ligases reported to be involved in bacterial ubiquitination are also implicated in crucial cellular functions (*6, 10*) and K48-Ub chains act as a major signal for cell’s own proteostatis, we hypothesized that similar principles could be adopted by the host for identification of K48-Ub substrate on bacterial surface. Since, for host proteins, presence of a tripartite motif (a primary degron sequence followed by a proximal lysine residue and a disordered region in-between) is reported to be a pre-requisite for K48-ubiquitination (*11*), we screened the surface proteins of SPN for the presence of similar features (Fig.1G). Our search through the pool of SPN proteins suggested BgaA and PspA to be the putative targets for ubiquitination on its surface (Fig.S1A,B). BgaA is a β-galactosidase which is reported to function as an adhesin for SPN, while PspA is a choline-binding protein that binds lactoferrin and is required for complement evasion (*12, 13*). Indeed, a drastic reduction (∼50%) was observed in association of K48-Ub chain type for both Δ*bgaA* and Δ*pspA* mutants without any change in K63-Ub levels (Fig.1H). The validity of our predicted targets was further proved by complementation and strengthened by utilizing Δ*hysA* (SPN surface protein which does not fulfill tripartite degron criteria) mutant as a control to score K48-Ub association levels (Fig.1H). We further explored the effect of K48-Ub decoration on bacterial clearance. Expectedly, absence of K48-Ub substrates impeded bacterial clearance resulting in significantly improved intracellular persistence for both mutant SPN strains (∼1.8 fold for Δ*bgaA* and ∼2 fold for Δ*pspA*) (Fig.1J,K). These effects were pronounced in double knock-out strain (Δ*bgaA*Δ*pspA*), suggesting the non-redundant nature of these ubiquitin substrates (Fig.1H,L). The universality of our substrate prediction approach was further validated by identification of an outer membrane protein, RlpA on STm, as a putative substrate for ubiquitination (Fig.1G). Indeed, the Δ*rlpA* mutant exhibited ∼1.5 fold reduced association with K48-Ub compared to WT STm (Fig.1I), establishing the ubiquitous applicability of our strategy. To the best of our knowledge, these are the first bacterial surface proteins reported to be recognized by host ubiquitination machinery for pathogen sensing and clearance.

**Fig. 1.**
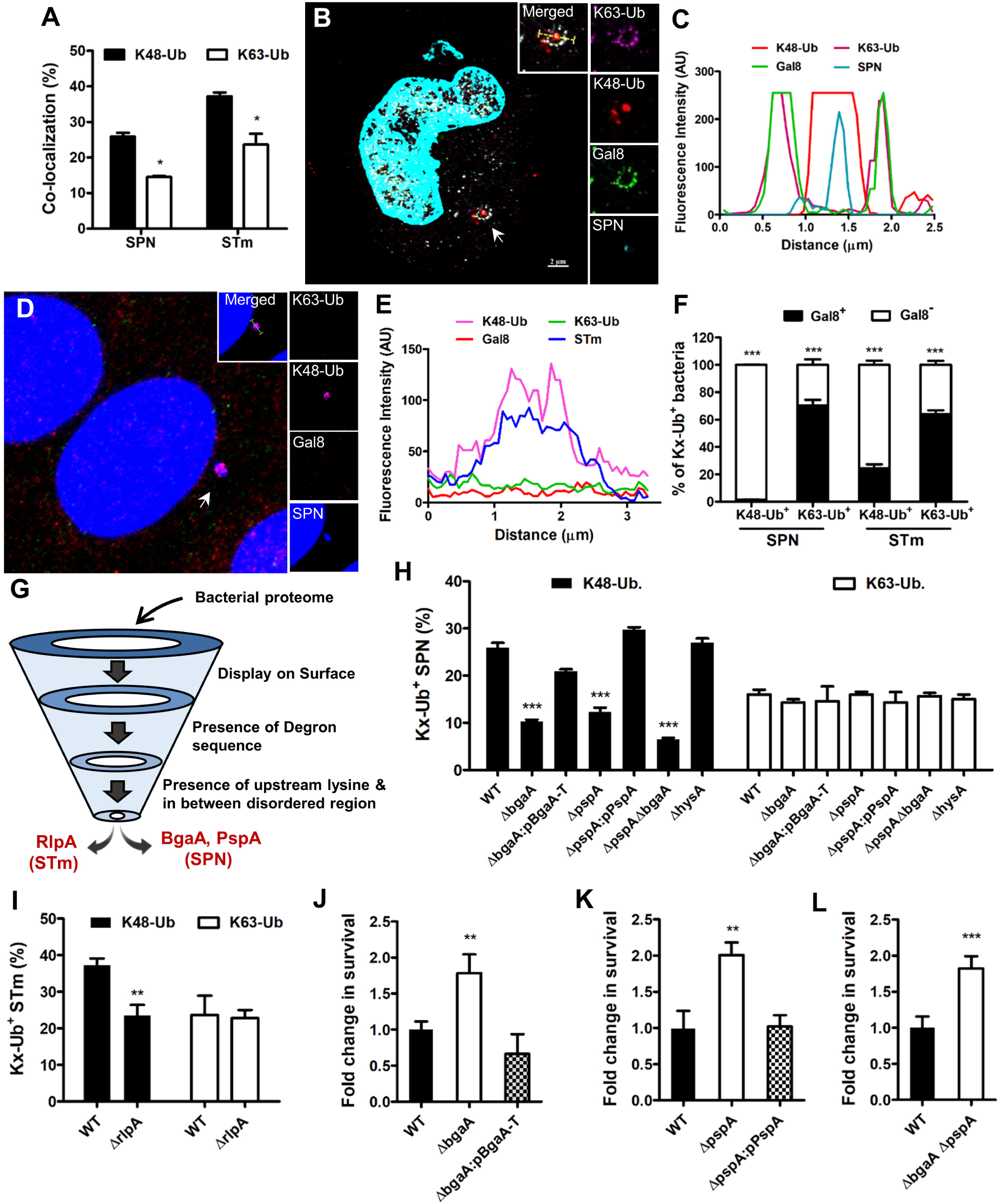
Identification of bacterial surface proteins as substrates for ubiquitination. **A**. Percentage of co-localization of SPN and STm with K48-Ub or K63-Ub at 9 h post infection in A549 and 3 h post infection in HeLa cells, respectively. n>100 bacteria/coverslip. **B**. Structured Illumination Micrograph of A549 cells showing association of SPN (Cyan) with K48-Ub (Red) inside a damaged bacteria containing endosome marked with Gal8 (Green) and K63-Ub (Magenta) at 9 h post infection. Arrowhead designate event shown in inset. Scale bar, 2 μm. **C**. Fluorescent line-scan across the yellow line in the merged inset in “**B**”. **D**. Confocal micrograph showing decoration of cytosolic STm (blue) with K48-Ub (magenta), devoid of K63-Ub (red) or Gal8 (green) in HeLa cells 3 h post infection. Arrowhead depicts event shown in inset. Scale bar, 5 µm. **E**. Fluorescent line-scan across the yellow line in the merged inset in “**D**”. **F**. Percentage of cytosolic (Gal8^-^) or damaged vacuole bound (Gal8^+^) SPN or STm marked with K48 and K63-Ub chain types. n>50 bacteria/coverslip. **G**. Schematic of screening for putative K48-Ub target proteins in bacteria. **H-I**. Comparison of association of K48 or K63-Ub with different SPN (**H**) and STm (**I**) strains following 9 h post infection in A549 cells or 3 h post infection in HeLa cells, respectively. n>100 bacteria/coverslip. **J-L**. Fold change in intracellular survival of different SPN strains in A549 cells normalized to WT SPN at 9 h post infection. Statistical significance was assessed by two way ANOVA (Bonferroni test) (A, F), one-way ANOVA (Dunnett’s test) (H, I, J, K) and two-tailed unpaired student’s t-test (L). *P < 0.05, **P < 0.01, ***P < 0.005. Data are mean ± SD of 3 independent biological replicates.

Following substrate identification, we aimed to test the critical features of the tripartite motif which form the backbone of our screen (Fig.2A, Fig.S2A). Deletion of the degron sequence (^102^VTPKEE^107^) in BgaA resulted in ∼50% drop in association of K48-Ub chain type compared to WT SPN (Fig.2C,D). Intriguingly, this reduction was comparable to the Δ*bgaA* knock-out strain strengthening the degron-specific phenotype. Apart from the degron sequence, presence of a lysine residue in close vicinity is crucial for attachment of the ubiquitin moiety to the substrate. Since in BgaA, the degron sequence is accompanied by two proximal lysines (K96, K97), we mutated both lysine residues (K96R, K97R), which resulted in 62% reduction in K48-ubiquitination compared to WT SPN (Fig.2D). Interestingly, among them, only K97R substitution led to a significant decrease (∼68%) in K48-ubiquitination, while K96R substitution resulted in only ∼31% reduction (Fig.2D). This result pinpointed to a precise mechanism adopted by host E3 ligases to select a particular lysine residue for tagging ubiquitin substrates. Expectedly, deletion of degron sequence and mutation of the marked lysine residue (K97) inhibited SPN clearance, leading to 1.5-2 fold increase in bacterial persistence ability (Fig.2E). Similar to BgaA, degron sequence (^327^PETPAPE^333^) deletion and lysine mutation (K315R) in PspA also exhibited significantly reduced K48-Ub association coupled with prolonged surviving ability (Fig.S2C,D).

**Fig. 2.**
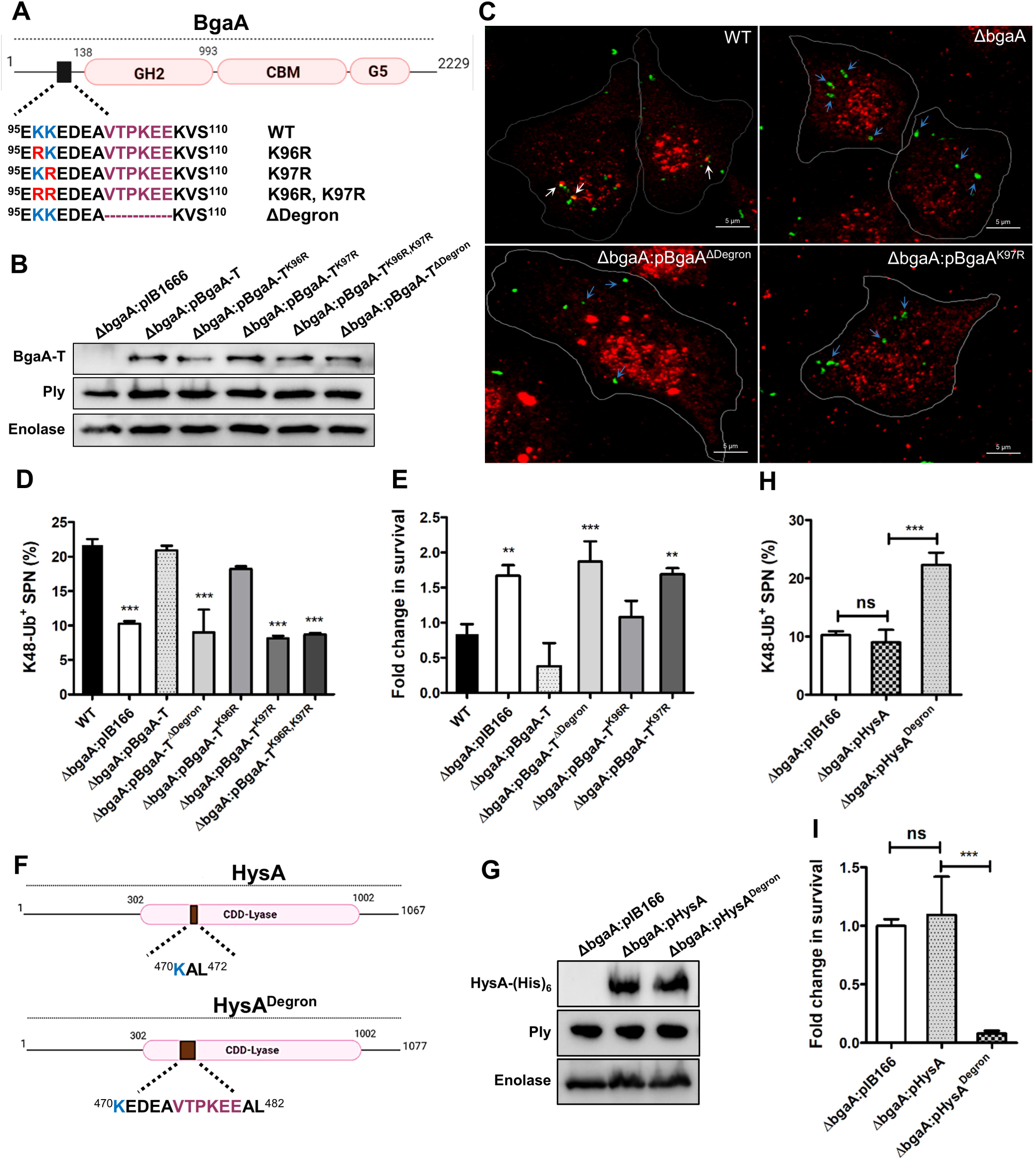
Significance of degron motif and proximal lysine in ubiquitination of BgaA. **A**. Schematic diagram of BgaA depicting presence of degron and proximal lysine residues. Also shown are different mutant BgaA variants used in the study. **B**. Immunoblot showing expression of BgaA variants in Δ*bgaA* strains. Similar levels of pneumolysin (Ply) across strains ensure equal damage to endosome membrane. Enolase was used as a loading control. **C**. Confocal micrographs of A549 cells at 9 post infection showing decoration of different SPN strains with K48-Ub. White arrowheads designate SPN associated with K48-Ub and blue arrowhead depicts absence of co-localization. Scale bar, 5 μm. **D**. Percentage of association of K48-Ub chain with different SPN strains at 9 h post infection in A549 cells. Δ*bgaA*:pIB166 designates vector control. n>100 bacteria/coverslip. **E**. Fold change in intracellular survival of different SPN strains in A549s normalized to WT at 9 h post infection. **F**. Schematic diagram of a non-ubiquitinable target protein HysA and its variant following incorporation of degron sequence of BgaA. **G**. Western blot showing expression of HysA and its variant in Δ*bgaA* background. Expression of Ply was similar across the strains and Enolase was used as loading control. **H**. Percentage of association of K48-Ub with different SPN strains carrying variants of HysA at 9 h post infection in A549 cells. n>100 bacteria/coverslip. **I**. Fold change in intracellular persistence ability of different SPN strains harboring HysA variants in A549s normalized to Δ*bgaA* at 9 h post infection. Statistical significance was assessed by one-way ANOVA followed by Dunnett’s test (D, E) or Tukey’s test (H, I). **P < 0.01, ***P < 0.005. Data are mean ± SD of 3 independent biological replicates.

To unequivocally prove our concept of K48-Ub substrate selection, we next engineered the SPN surface protein HysA, lacking a primary degron sequence (Fig.2F,G; Fig.S1C-E). We hypothesized that addition of degron sequence (in a structurally disordered region containing a lysine residue) will trigger K48-ubiquitination of the non-ubiquitinable HysA protein. Indeed, K48-Ub association levels of SPN strain carrying the engineered HysA protein (Δ*bgaA*:pHysA^Degron^) was found to be significantly higher compared to either Δ*bgaA* or Δ*bgaA*:pHysA strains (Fig.2H). The enhanced decoration of Δ*bgaA*:pHysA^Degron^ strain with K48-Ub, robbed the survival gain provided by the absence of degron in Δ*bgaA* or Δ*bgaA*:pHysA (Fig.2I). A similar phenotype was also observed when HysA protein was engineered with addition of degron sequence present in PspA (Fig.S2F,G). Importantly, all the engineered SPN strains produced similar levels of the pore forming toxin, pneumolysin, which is prerequisite for endomembrane damage and subsequent ubiquitination (Fig.2B,G; Fig.S2B,E). This nullifies the possible contribution of low or extensive membrane damage promoting dramatic change of ubiquitination levels in mutant SPN strains. Collectively, these suggest that artificial addition of degron sequence would promote ubiquitin mediated detection and elimination of pathogens that do not follow same suit.

The canonical degron sequence present in BgaA as well as PspA is predicted to be identified by the SCF^FBW7^ E3 ligase (*11*), which is involved in regulation of cell cycle and growth (14) and composed of two conserved proteins Skp1 and a member of Cullin family protein in addition to variable F-box element that provides substrate specificity (*15*). To verify involvement of SCF^FBW7^ in SPN ubiquitination, we first assessed association of FBXW7 with SPN. Association of ∼31% of intracellular SPN with FBXW7 (Fig.3A,B), which is similar to K48-Ub levels, confirms our assumption. To strengthen our findings, marking of bacteria with K48-Ub chains was examined by immunofluorescence following downregulating the expression of Cullin1, Skp1, and FBXW7 genes using targeted siRNAs (Fig.3C). We observed significant reduction in SPN association with K48-Ub in Cullin1, Skp1, and FBXW7 knock-down cells (Fig.3D-F) which in turn triggered ∼1.5 fold increase in the SPN persistence inside host cells (Fig.3G-I). We further performed *in vitro* ubiquitination with purified BgaA-T (Fig.S3) and the SCF complex components to unambiguously demonstrate the role of SCF^FBW7^ E3 ligase in ubiquitination of BgaA. Our results showcased that recombinant SCF^FBW7^ ubiquitinated purified BgaA-T (Fig.3J), which confirms SCF^FBW7^ as the bonafide E3 ubiquitin ligase for BgaA. Critically, SCF^FBW7^ failed to ubiquitinate the degron deleted variant BgaA-T^ΔDegron^ as well as BgaA-T^K97R^ (Fig.3J), the lysine residue mutant tipped for ubiquitin attachment. These undoubtedly prove the novel role of SCF^FBW7^ E3 ligase in detection of cytosol dwelling pathogens and targeting them towards killing pathways. Since most target substrates of SCF complex requires phosphorylation for interaction with F-box protein, we tested the likelihood and impact of phosphorylation of bacterial substrates on K48-Ub coating of the pathogen. Bioinformatics analysis suggested that the degron sequence in BgaA involved a putative phosphorylable threonine residue (^102^VT*PKEE^107^). Indeed, we observed that SPN strain harboring BgaA-T^T103A^ mutation (Δ*bgaA*:pBgaA-T^T103A^) manifested ∼50% reduced co-localization with K48-Ub compared to WT (Fig.4A), revealing the relevance of phosphorylation in substrate recognition by SCF complex. Significantly, the decreased propensity of BgaA phosphorylation in Δ*bgaA*:pBgaA-T^T103A^ abrogated host’s ability to eliminate intracellular bacterial load (Fig.4B). In parallel to BgaA, PspA (Δ*pspA*:pPspA^T329A^) degron variant also showed a similar drop in K48-Ub co-localization and manifested prolonged intracellular persistence ability (Fig.S4). In general, targets (of SCF complex containing FBXW7 as a component) possessing threonine/serine (T/S*) next to a proline residue are phosphorylated by a proline-directed protein kinase, GSK3β (*16-18*). We therefore attempted to unravel the involvement of GSK3β in augmenting the substrate recognition. In an *in vitro* kinase assay, we observed that GSK3β could phosphorylate recombinant BgaA-T while BgaA-T^T103A^ variant remained non-phosphorylated (Fig.4C). This validated our earlier findings about the identity of the threonine residue in BgaA-T, the target for GSK3β mediated phosphorylation. As expected, targeted knockdown of GSK3β by siRNA (Fig.4D) led to ∼58% reduction in K48-ubiquitination of SPN (Fig.4E). This reduced ubiquitination following downregulation of GSK3β expression, was further reflected in diminished ability of the host to clear cell invaded pathogens (Fig.4F). Collectively, this provides the first evidence of a host kinase, specifically GSK3β, regulating ubiquitination of bacterial surface proteins for efficient clearance of pathogens (Fig.4G).

**Fig. 3.**
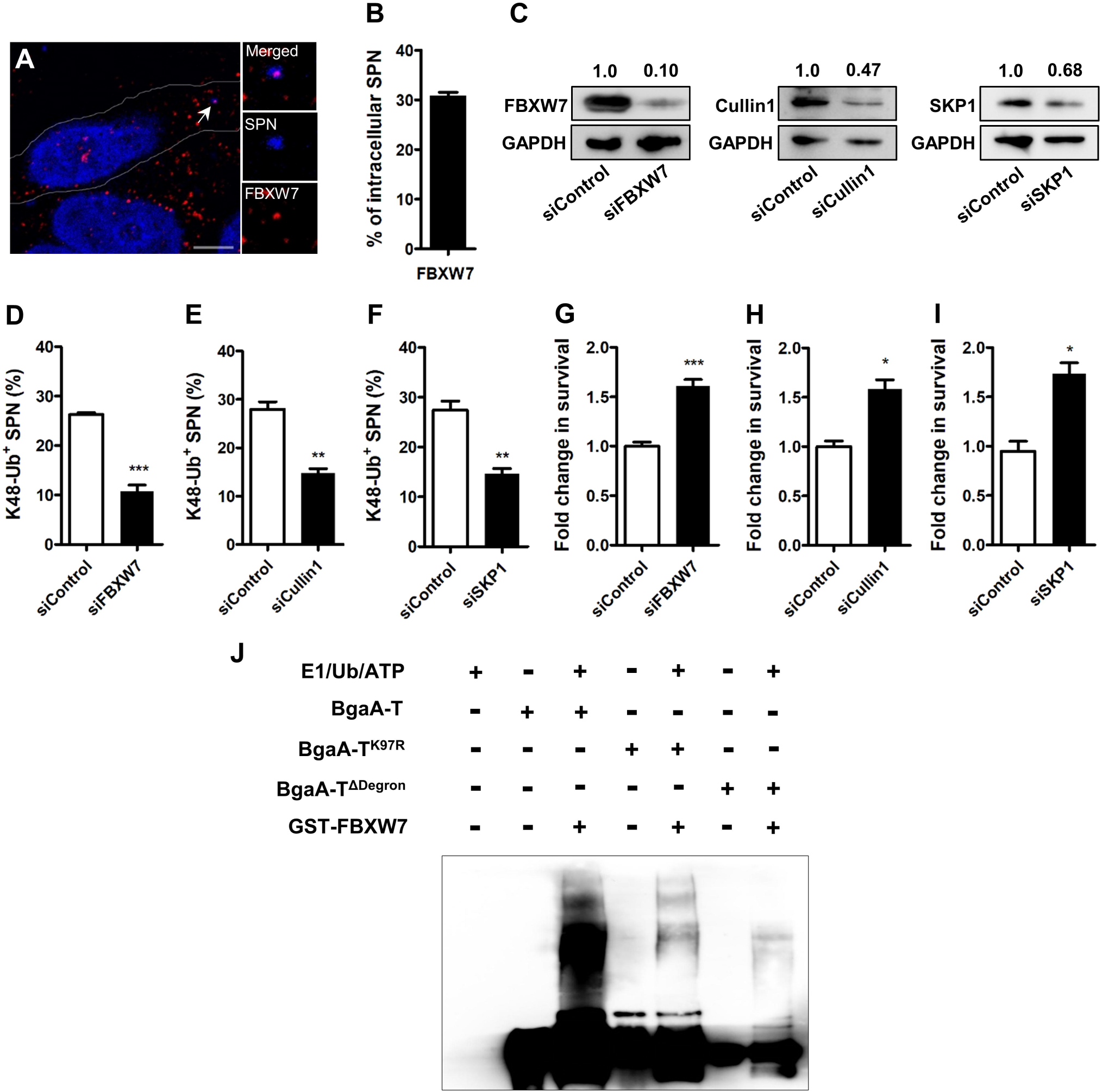
SCF^FBW7^ ubiquitinates surface proteins of cytosolic SPN. **A**. Confocal micrograph of A549 cells at 9 post infection showing association of SPN (blue) with FBXW7 (magenta), the substrate recognizing component of SCF^FBW7^ E3-ligase. Arrowheads designate event shown in inset. Scale bar, 5 μm. **B**. Percentage of association of SPN with FBXW7 in A549s at 9 h post infection. n>100 bacteria/coverslip. **C**. Immunoblot demonstrating knock-down in expression of FBXW7, Cullin1, SKP1, the key components of SCF^FBW7^ E3-ligase in A549 cultures following transfection with siFBXW7, siCullin1, and siSKP1, respectively. GAPDH served as loading control. Fold change in expression of individual targets relative to siControl (post normalization with GAPDH) is mentioned above the blot. **D-F**. Percentage of decoration of SPN with K48-Ub following knock down of FBXW7 (**D)**, Cullin1 (**E**) and Skp1 (**F**) in comparison to scrambled siRNA (siControl) transfected cells. n>100 bacteria/coverslip. **G-I**. Fold change in intracellular survival of SPN in A549 cells transfected with siFBXW7 (**G**), siCullin1 (**H**) and siSKP1 (**I**) and normalized to siControl. **J**. *In vitro* ubiquitination reaction demonstrating ubiquitination of BgaA-T, BgaA-T^K97R^ and BgaA-T^ΔDegron^. Recombinant active components of SCF complex (E1, E2(Cdc34), Rbx1, FBXW7) were incubated with purified BgaA-T-His_6_ variants and ubiquitin, ATP. Anti-BgaA antibody was used for immunoblotting. Statistical significance was assessed by two-tailed unpaired student’s t-test (D - I). *P < 0.05, **P < 0.01, ***P < 0.005. Data are mean ± SD of 3 independent biological replicates.

**Fig. 4.**
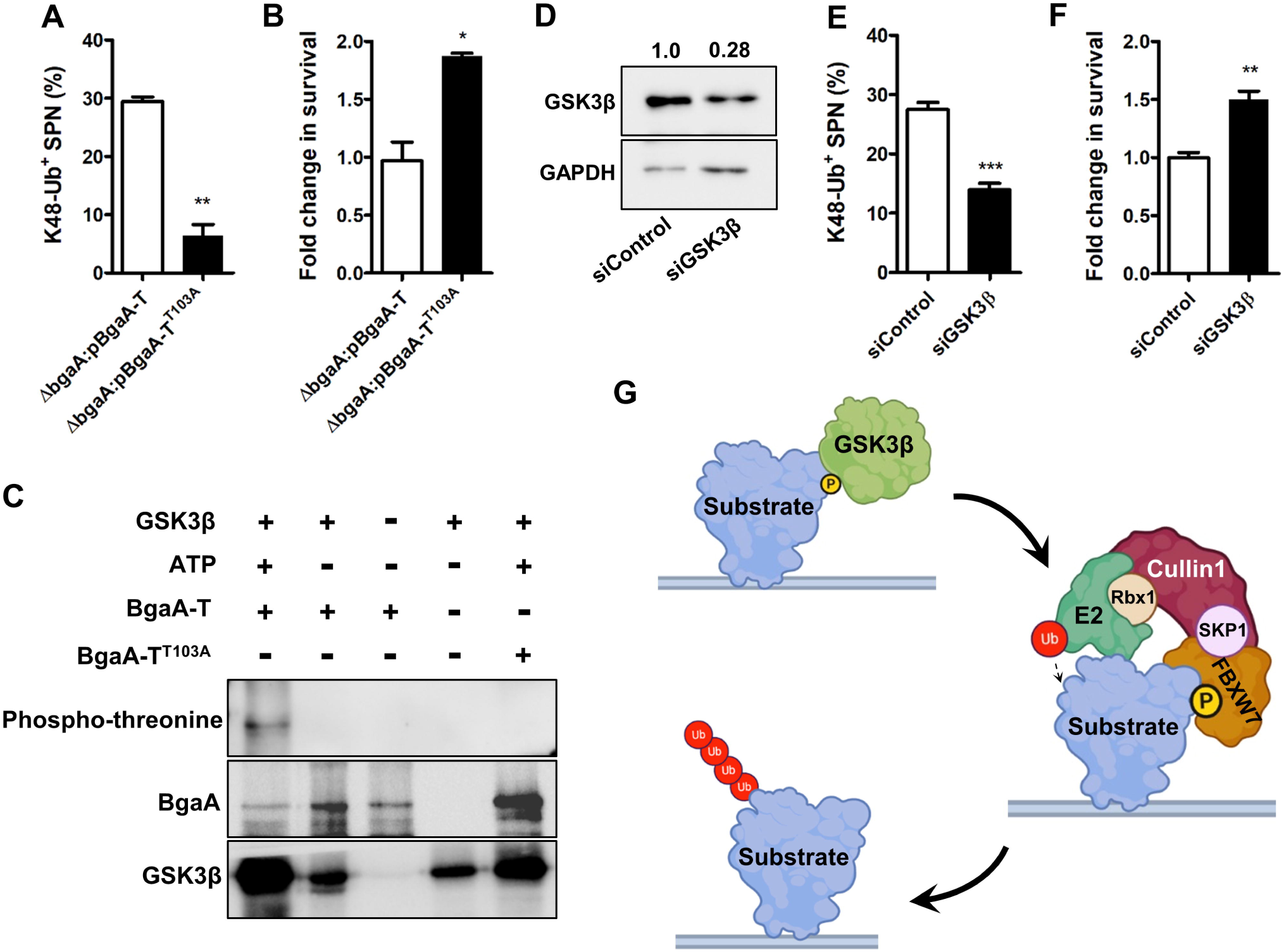
GSK3β mediated phosphorylation is key for ubiquitination of SPN surface proteins. **A**. Percent K48-Ub association with different SPN strains in A549 cells at 9 h post infection. n>100 bacteria/coverslip. **B**. Fold change in intracellular persistence ability of different SPN strains in A549s normalized to WT SPN at 9 h post infection. **C**. *In vitro* kinase assay assessing phosphorylation of Bga-T and Bga-T^T103A^ by recombinant GSK3β. Phosphorylated proteins were detected with anti-phospho threonine antibody. **D**. WB demonstrating knock-down in expression of GSK3β in A549 cultures following transfection with siGSK3β. GAPDH served as loading control. Fold change in expression of GSK3β with respect to siControl (post normalization with GAPDH) is mentioned above the blot. **E**. Percentage of association of SPN with K48-Ub following knock down of GSK3β. Cells transfected with scrambled siRNA (siControl) served as a control. n>100 bacteria/coverslip. **F**. Fold change in intracellular survival of SPN in A549 cells transfected with siGSK3β and normalized to siControl. **G**. Schematic depicting phosphorylation of degron motif in bacterial surface protein by GSK3β, followed by recruitment and ubiquitination by SCF^FBW7^ E3 ligase. Scheme was designed with help of Biorender® software. Statistical significance was assessed by two-tailed unpaired student’s t-test (A, B, E, F). *P < 0.05, **P < 0.01, ***P < 0.005. Data are mean ± SD of 3 independent biological replicates.

Metazoans employ ubiquitination as a versatile mechanism to maintain the sanctity of the cytosol, both from damaged proteins/organelles as well as from invading pathogens (*19, 20*). Given that it encounters variety of pathogens and diverse set of damaged proteins, identification of common motifs for substrate recognition and subsequent ubiquitination could be a smart strategy for resource optimization. In general, the proteins in eukaryotic cells destined for proteolysis are identified by tripartite degron motif (*11*). This prompted us to explore whether similar motifs could be found on pathogenic surface as a proxy for substrate recognition. Here, we demonstrated for the first time that host ubiquitin ligases utilize similar features to sense phylogenetically distinct pathogens. Pathogen recognition via common molecular pattern/motif (PAMPs) is a well characterized framework of innate immunity (*21*). Our results suggest that degron-like sequences in bacterial surface proteins act equivalent to PAMPs to impart intracellular pathogen surveillance. This modus operando could be further exploited by transforming a non-substrate protein into an active ubiquitination substrate for potent sensing and clearance of a pathogen. The fail-safe mode of operation offered by this strategy becomes more apparent as the mutant bacterial strains still exhibited substantial residual ubiquitination, highlighting utilization of other substrates by ubiquitin machinery for robust pathogen interception. A similar pathogen sensing mechanism involving Guanylate binding proteins (GBPs) is also implicated in bacterial clearance; however, unlike ubiquitination their role is still reported to be restricted to selective category of pathogens (*22, 23*). Interestingly, few pathogens are known to actively modify surface proteins, protecting them from ubiquitination and subsequent killing, thus emphasizing the significance of surface localization of the ubiquitin substrate (*4*). Additionally, multiple obligate intracellular pathogens have evolved strategies to de-ubiquitinate themselves or host regulatory components to evade ubiquitin mediated clearance (*24*). Taken together, these underscore ubiquitin-mediated alarm arousal as a fundamental intracellular pathogen sensing mechanism central to host defenses.

Our study further demonstrated the central role of SCF^FBW7^ E3 ligase in bacterial ubiquitination augmenting pathogen degradation. This implies that individuals carrying a loss of function mutations in SCF complex genes could be highly susceptible to bacterial infections. Indeed, a recent cohort-based study indicated that almost 43% of the CLL (Chronic Lymphocytic Leukaemia) patients, caused by mutations in the FBXW7 gene, succumb to bacterial pneumonia and sepsis followed by fungal infections (*25*). Our findings therefore provide an unexpected molecular explanation of the enhanced risk of infections of the CLL patients, especially to respiratory pathogens. The link between genetic polymorphism in E3 ligase genes and susceptibility to bacterial infections is also substantiated by Parkinson’s patients, who are vulnerable to typhoid fever or leprosy (*10*), suggesting significant contribution of E3 ligases in host immunity against bacterial infections and maintenance of cellular steady-state. In conclusion, we deciphered a universal language of sensing cytosol dwelling pathogens that could be efficiently applied by the host to recognize any microorganism for subsequent elimination. Collectively, these shed light on the understanding of the rudimentary cellular immune processes, which could be harnessed to intensify the antibacterial immunity.

## Materials and Methods

### Cell culture

Human lung alveolar carcinoma (type II pneumocyte) cell line A549 (ATCC No. CCL-185) and cervical adenocarcinoma cell line HeLa (ATCC No. CRM-CCL-2), both were cultured in DMEM (HiMedia) supplemented with 10% fetal bovine serum (FBS, Gibco) at 37°C and 5% CO_2_.

### Bacterial strains and growth conditions

*Streptococcus pneumoniae* (SPN) (R6, serotype 2, gift from Prof. TJ Mitchell, Univ. of Birmingham, UK) was grown in Todd-Hewitt broth (THB) supplemented with 1.5% yeast extract at 37□C in 5% CO_2_. The following antibiotics were used to grow SPN cultures when required: kanamycin (200 µg/ml), spectinomycin (100 µg/ml) and chloramphenicol (4.5 µg/ml). 900 µl of 0.4 OD600 grown SPN culture was mixed with 600 µl of 80% sterile glycerol (32% final glycerol concentration) and stored in -80□C deep freezer. These glycerol stocks were used as starting inoculum for all experiments.

*E. coli* DH5α and *Salmonella* Typhimurium (STm, ATCC 14028) cultures were grown in Luria-Bertani broth (LB) at 37□C under shaking conditions (200 rpm) and when necessary following antibiotics were used: kanamycin (50 µg/ml), spectinomycin (100 µg/ml), chloramphenicol (20 µg/ml) and ampicillin (100 µg/ml). For all infection assays, bacteria were grown till OD_600_ ∼ 0.4, followed by resuspension in PBS to similar density before infecting the host cells.

### Screening of putative ubiquitin target proteins

All surface proteins present in the STm and SPN proteome were first identified from published literature. Complete protein sequences of the surface proteins were obtained from Uniprot, which were then used to identify presence of degron motifs. A set of 29 putative degron motifs were curated from published literature (*11, 26, 27*) and the Eukaryotic Linear Motif database. Python’s regex module was used to identify the location of all such curated motifs in the surface protein sequences. A linear search was further performed to identify the presence of a lysine residue in close proximity (8-14 amino acid residues) to every identified degron motif. This lysine residue is presumed to act as the attachment site for ubiquitin moiety. Finally, the shortlisted protein sequences were fed into IUPred (*28*) a protein structure predicting tool, to locate the presence of disordered region in between the degron motif and proximal lysine. The resultant proteins (BgaA and PspA for SPN; RlpA for STm) were selected for evaluation as ubiquitination targets and decoding their role in pathogen clearance.

### Bacterial strain construction

Allelic exchange by homologous recombination was carried out using gene flanking regions carrying an inserted antibiotic cassette for generation of mutant strains in both SPN and STm (Table S1). For SPN, 500 bp upstream and downstream regions of *bgaA, pspA* and *hysA* genes were amplified from the genome with appropriate primers (Table S2) and assembled in pBKS vector. Following cloning of these fragments, antibiotic resistance cassette (Spectinomycin for *bgaA* as well as *pspA* and Chloramphenicol for *hysA*) was inserted into this construct. The linearized recombinant plasmids were then transformed into WT SPN using competence stimulating peptide 1 (CSP-1) (GenPro Biotech), recombinants were selected using respective antibiotic and gene replacement was confirmed by PCR as well as sequencing of the respective gene loci. For STm, λ-red recombinase method was used for generation of gene-deletion mutants (*29*). Briefly, sequences homologous to the ends of the *rlpA* gene were appended in the primer sequences that are used for amplification of the kanamycin resistance cassette. The cassette was then electroporated into the WT STm strain and knockouts generated by homologous recombination were selected based on kanamycin resistance. Gene deletion was confirmed using PCR and sequencing of the gene locus. Full length *pspA, hysA* and a trauncated *bgaA-T* (1-3168bp) were cloned under P23 promoter in the shuttle vector pIB166(*30*) and used for complementation. These recombinant plasmids were also used for site directed mutagenesis to generate different variants of *bgaA-T* (*bgaA-T*^*K96R*^, *bgaA-T*^*K97R*^, *bgaA-T*^*K96R,K97R*^ and *bgaA-T*^Δ*Degron*^), *pspA* (*pspA*^*K314R*^, *pspA*^*K315R*^, *pspA*^*K314R,K315R*^ and *pspA*^Δ*Degron*^) and *hysA* (*hysA*^*Degron*^) using appropriate primer sets (Table S2) and transformed into Δ*bgaA* and Δ*pspA* mutants. All clones were verified by DNA sequencing. Expression of different variants of BgaA, PspA and HysA were confirmed by western blot using appropriate antibodies.

### Antibodies and reagents

Anti-Enolase and anti-PspA serum (Prof. S. Hammerschmidt, University of Greifswald, Germany); Anti-BgaA serum (Dr. Samantha King, The Ohio State Univ., USA). Antibodies specific for His_6_ (Invitrogen MA1-21315), K48-Ub linkage (Millipore 05-1307), K63-Ub linkage (Millipore 14-6077-80), Pneumolysin (Santacruz sc-80500), FBXW7 (Bethyl A301-721A), SKP1 (Invitrogen MA5-15928), Cullin1 (Invitrogen 71-8700), GAPDH (Millipore MAB374), GSK3β (CST D5C5Z), Phosphothreonine (CST 9381S), TEPC-15 (Sigma M1421) were procured. The following secondary antibodies were used: HRP tagged anti-rabbit (Biolegend 406401), HRP tagged anti-mouse (Biolegend 405306), Anti-rabbit Alexa Fluor 488 (Invitrogen A27206), anti-rabbit Alexa Fluor 555 (Invitrogen A31572), Biotin conjugated anti-mouse IgaA (Life Tech. M31115), anti-mouse Alexa Fluor (Invitrogen A31570), anti-mouse Alexa Fluor 488 (Invitrogen A21202), anti-goat Alexa Fluor 633 (Invitrogen A21082).

### Protein expression and purification

BgaA-T and its variants were cloned in pET28 using XbaI/NotI restriction sites. Recombinant plasmids encoding BgaA-T with N-terminal His-tag were transformed into *E. coli* BL21 (DE3) cells for protein expression. Freshly transformed colonies were grown in LB containing 50 μg/ml kanamycin at 37°C on a shaker incubator for 12 h. 1% of the primary culture was added to l L of LB broth and incubated at 37°C on a shaker incubator till the OD_600nm_ reached between 0.6–0.8. Protein expression was induced by the addition of 100 μM isopropyl-1-thiogalactopyranoside (IPTG) and growing the culture further at 37°C for 5 - 6 h with agitation at 150 rpm. The cells were harvested by centrifugation at 6,000 rpm for 10 min at 4°C. The cell pellet was resuspended in buffer A (25 mM Tris, pH 8.0 and 300 mM NaCl) and lysed by sonication. Cell debris was separated by centrifugation (14,000 rpm, 50 min, 4°C) and supernatant was applied on to a Ni-NTA column, equilibrated with buffer A. The column was washed with 10 column volumes of buffer A and the His-tagged proteins were eluted with imidazole (250 mM) in buffer A. All the variants of BgaA-T were expressed and purified using the same procedure. FBXW7 was subcloned in pGEX-4T-1 vector having an N-terminal GST tag from pCMV6-Entry-FBXW7 vector using EcoRI restriction enzyme. GST-tagged FBXW7 was expressed in *E. coli* BL21(DE3) cells following induction of protein expression with 0.1 mM IPTG and growth at 30°C for 4 h. GST-FBXW7 from *E. coli* crude extract was purified using Glutathione-Sepharose (GE Healthsciences) column chromatography. Fractions containing purified proteins were pooled and concentrated up to 0.5 mg/ml using 10 kDa molecular weight cut off filter (Amicon) by centrifugation at 4,700 rpm at 4°C. Purity of the proteins was checked on SDS-PAGE followed by staining with Coomassie blue.

### siRNA directed gene knockdown

For RNAi mediated knock-down of specific genes, the following siRNAs were used: siFBW7 (Dharmacon ON-TARGET SMARTpool L-004246-00-0005); siCullin1 (Qiagen 1027423); siSKP1 (Qiagen SI00301819) and siGSK3β (Dharmacon ON-TARGET SMARTpool L-003010-00-0005). Briefly, A549 cells grown in 24 well plate were transiently transfected with gene specific siRNAs or scramble (siControl) (150 pmol) using Lipofectamine 3000 as per manufacturer’s instructions. 36 h post transfection, cells were processed for western blotting as well as immunofluorescence or penicillin-gentamycin protection assays.

### Structure prediction and modelling

All structures were predicted by Phyre^2^ (Protein Homology/analogY Recognition Engine V 2.0) (*31*) and visualized using PyMOL (The PyMOL Molecular Graphics System, Version 2.0 Schrödinger, LLC). Structures were color coded in PyMol based on IUPred(*28*) scores ranging from ordered (blue) to disordered (red) through white.

### Western blotting

SPN cultures grown to 0.4 OD_600nm_ were lysed by sonication and crude extracts were collected following centrifugation (15000 rpm, 30 min, 4°C). For A549s, monolayers were washed several times with PBS and lysed in ice-cold RIPA buffer (50 mM Tris-Cl, pH 7.89, 150 mM NaCl, 1% Triton X-100, 0.5% Sodium deoyxycholate, 1% SDS) containing protease inhibitor cocktail (Promega), Sodium fluoride (10 mM) and EDTA (5 mM). The cell suspension was briefly sonicated and centrifuged to collect cell lysates. Proteins present in bacterial or A549 cell lysates (10 μg or 20 μg) were separated on 12% SDS-PAGE gels and transferred to activated PVDF membrane. Following blocking in 5% skimmed milk the membranes were probed with appropriate primary and HRP-tagged secondary antibodies. The blots were finally developed using ECL substrate (Bio-Rad).

### Penicillin-gentamicin protection assay

SPN strains grown until OD_600nm_ 0.4 in THY were pelleted, re-suspended in PBS (pH 7.4) and diluted in assay medium for infection of A549 monolayers with multiplicity of infection (MOI) of 10. Following 1 h of infection, the monolayers were washed with DMEM and incubated with assay medium containing penicillin (10 µg/ml) and gentamicin (400 µg/ml) for 2 h to kill extracellular SPN. Cells were then lysed with 0.025% Triton X-100 and the lysate was plated on Brain Heart Infusion agar plates to enumerate viable SPN. Percentage invasion was calculated as (CFU in the lysate / CFU used for infection) × 100. To assess intracellular survival, at 8 h post infection (from the beginning of penicillin-gentamycin treatment) cell lysates were prepared as mentioned above, spread plated and surviving bacteria were enumerated. Survival efficiency (%) was represented as fold change in percent survival relative to control at indicated time point (normalized to 0 h).

### Immunofluorescence

For immunofluorescence assay, A549 or HeLa cells were grown on glass cover slips and infected with SPN or STm strains at MOI ∼ 25 for 1 h followed by antibiotic treatment for 2 h. At desired time points post infection (8 h for SPN infection in A549s and 3 h for infection with STm in HeLa), cells were washed with DMEM, and fixed with ice-chilled Methanol at -20°C for 10 min. Further, the coverslips were blocked with 3% BSA in PBS for 2 h at RT. Cells were then treated with appropriate primary antibody in 1% BSA in PBS for overnight at 4°C, washed with PBS and incubated with suitable secondary antibody in 1% BSA in PBS for 1 h at RT. Finally, coverslips were washed with PBS and mounted on glass slides along with VectaShield® with or without DAPI (Vector Laboratories) for visualization using a Laser Scanning Confocal microscope (LSM 780, Carl Zeiss) under 40X or 63X oil objectives. The images were acquired after optical sectioning and then processed using Zen lite software (Version 5.0.). Super resolution microscopy was performed similarly using Elyra 7 (Carl Zeiss) in Structured Illumination Microsocopy (SIM) mode. For co-localization analysis, bacteria were scored by visual counting of n >100 bacteria per replicate.

### *In vitro* ubiquitination

For *in vitro* ubiquitination assays, recombinant proteins were purified as described earlier (*32*). The SCF^FBW7^ E3 ligase complex was incubated with recombinant 0.1 mM E1 (UBE1; Boston Biochem), 0.25 mM E2 (cdc34; Boston Biochem), and 2.5 μg/ml ubiquitin (Boston Biochem) in the presence of purified BgaA-T and its variants. Ubiquitylation reactions were performed in assay buffer (50 mM Tris (pH 8), 5 mM MgCl_2_, 5 mM ATP, 1 mM β– mercaptoethanol, 0.1% Tween 20) for 2 h at 25°C. The reactions were stopped with 5× laemmli buffer, resolved on SDS-PAGE gels, and analyzed by immunoblotting using an anti-BgaA antibody.

### *In vitro* kinase assay

To perform *in vitro* kinase assay, GSK3β kinase enzyme system (Promega) was used as per manufactures protocol with following modifications. Briefly, 3 μg of recombinant BgaA-T was added to 0.5 μg of active GSK3β with 400 µM ATP in reaction buffer composed of 40mM Tris, 7.5; 20mM MgCl_2_; 0.1mg/ml BSA; 50μM DTT. The reaction was incubated for 2 h at 30□C and stopped by adding 1X Laemmli buffer. The samples were then boiled and electrophoresed for western blotting as mentioned previously. Phosphorylated BgaA-T was detected with anti-phosphothreonine Ab.

### Statistical analysis

GraphPad Prism version 5 was used for statistical analysis. Statistical tests undertaken for individual experiments are mentioned in the respective figure legends. *p*<0.05 was considered to be statistically significant. Data were tested for normality and to define the variance of each group tested. All multi-parameter analyses included corrections for multiple comparisons and data are presented as mean□±□standard deviation (SD) unless otherwise stated.

## Acknowledgements

We acknowledge the “Bio-safety Level 2 Facility” and “Confocal Microscopy Facility” at IIT Bombay. SA, KS and SS acknowledge financial support from IIT Bombay. SG and JJ, SB, SR acknowledges fellowship from CSIR and UGC, Govt. of India, respectively. AB acknowledges research funding Science and Engineering Research Board, Govt. of India (Grant No. SPR/2019/000808). The funders had no role in study design, data collection and analysis, decision to publish, or preparation of the manuscript

## Figure legends

**Fig. S1. Predicted structures of ubiquitination targets. A-C**. Structures of BgaA (**A**), PspA (**B**) and HysA (**C**) as predicted by Phyre^2^ and visualized using PyMOL. Shown also are zoomed in areas of the protein containing the degron sequence and the lysine residue tagged for ubiquitin attachment. **D-E**. Zoomed in version of predicted HysA structures following incorporation of the degron sequence of BgaA (**D**) or PspA (**E**) into HysA. Color coding is based on scores predicted by IUPred, ranging from ordered (blue) to disordered (red) through white where green depicts a degron region while lysine residue is shown in yellow ball and stick conformation. The scheme with respective IUPred scores and its color coding is given below.

**Fig. S2. PspA is also a target for ubiquitination. A**. Schematic diagram of PspA depicting presence of degron and proximal lysine residues. Also shown are different mutant PspA variants used in the study. **B**. Immunoblot showing expression of PspA variants in Δ*pspA* strains. All strains expressed similar levels of pneumolysin (Ply) ensuring equal damage to endosome membrane. Enolase was used as a loading control. **C**. Percentage of decoration of different SPN strains with K48-Ub at 9 h post infection in A549 cells. n>100 bacteria/coverslip. **D**. Fold change in intracellular survival of different SPN strains carrying mutations in the putative lysine residues or deletion of degron sequence, in A549s normalized to WT SPN at 9 h post infection. **E**. Western blot showing expression of HysA variant with degron sequence of PspA incorporated in a disordered region in Δ*pspA* background. Similar levels of pneumolysin (Ply) across strains ensure equal damage to endosome membrane. Enolase was used as a loading control. **F**. Percentage of association of K48-Ub with HysA variant expressing Δ*pspA* mutant strain at 9 h post infection in A549 cells. n>100 bacteria/coverslip. **G**. Fold change in intracellular persistence ability of Δ*pspA* mutant strain harboring degron motif added HysA variant in A549s normalized to Δ*pspA* at 9 h post infection. Statistical significance was assessed by one-way ANOVA followed by Dunnett’s test (C, F) or Tukey’s test (D, G). **P < 0.01, ***P<0.005. Data are mean ± SD of 3 independent biological replicates.

**Fig. S3. Purified BgaA variants**. SDS-PAGE analysis of different BgaA variants expressed as His-tagged protein and purified via Ni-NTA column chromatography for use in *in vitro* ubiquitination and *in vitro* kinase reaction. BgaA-T (**A**); BgaA-T^K97R^ (**B**); BgaA-T^ΔDegron^ (**C**); BgaA-T^T103A^ (**D**).

**Fig. S4. PspA is phosphorylated for subsequent ubiquitination. A-B**. Immunoblot analysis of PspA (**A**) and BgaA (**B**) complemented phosphodegron mutant variants **B**. Percentage of K48-Ub association with different SPN strains harboring PspA variants in A549 cells at 9 h post infection. n>100 bacteria/coverslip. **C**. Fold change in intracellular persistence ability of phosphorylable and non-phosphorylable PspA variant harboring SPN strains in A549s normalized to WT SPN at 9 h post infection. Statistical significance was assessed by two-tailed unpaired student’s t-test. **P < 0.01. Data are mean ± SD. of 3 independent biological replicates.

**Table S1**. List of bacterial strains.

**Table S2**. List of primers.

