## Supplementary figures and images for "A novel innate pathogen sensing strategy involving ubiquitination of bacterial surface proteins"

### Fig. S1

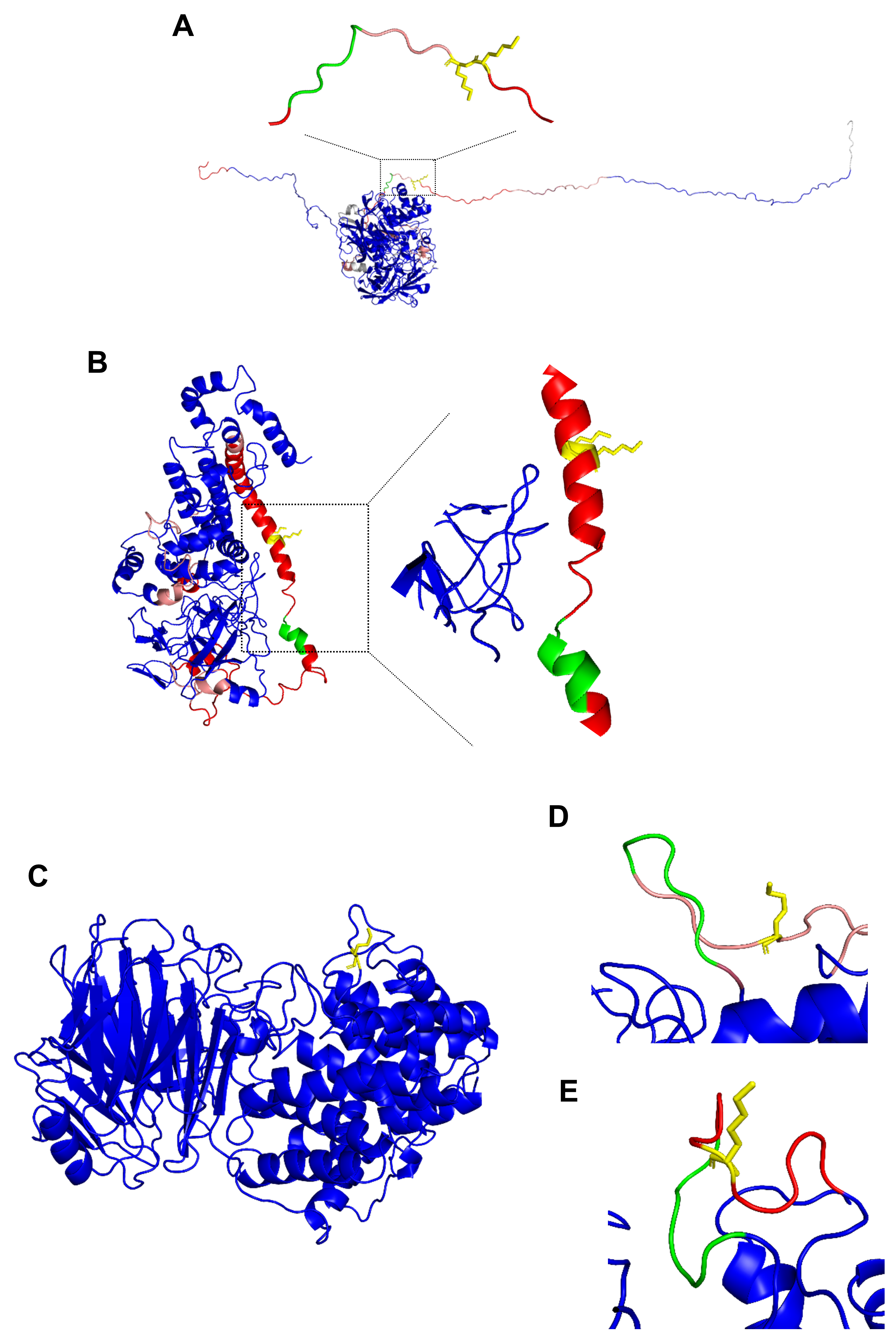

### Fig. S2

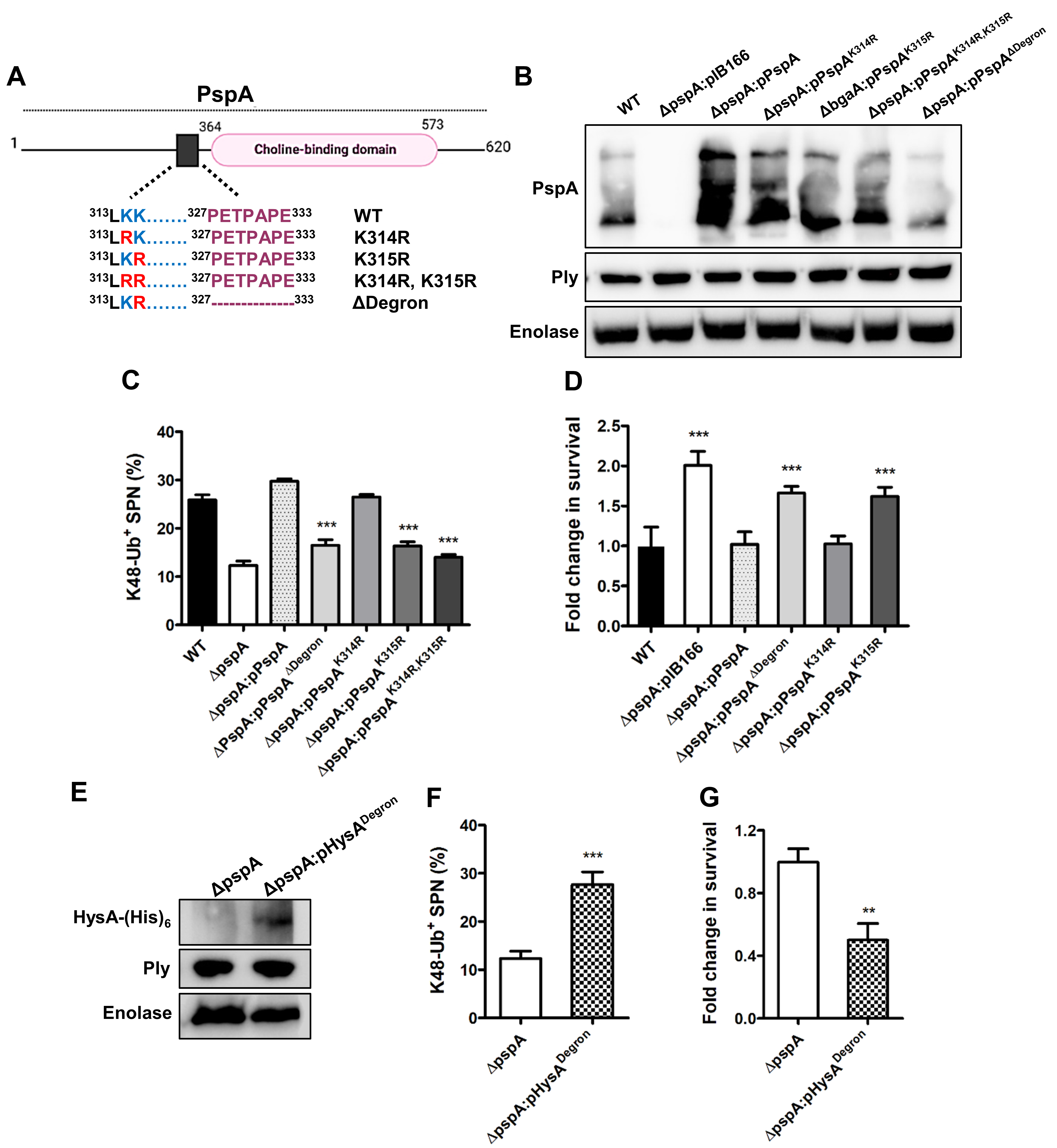

### Fig. S3

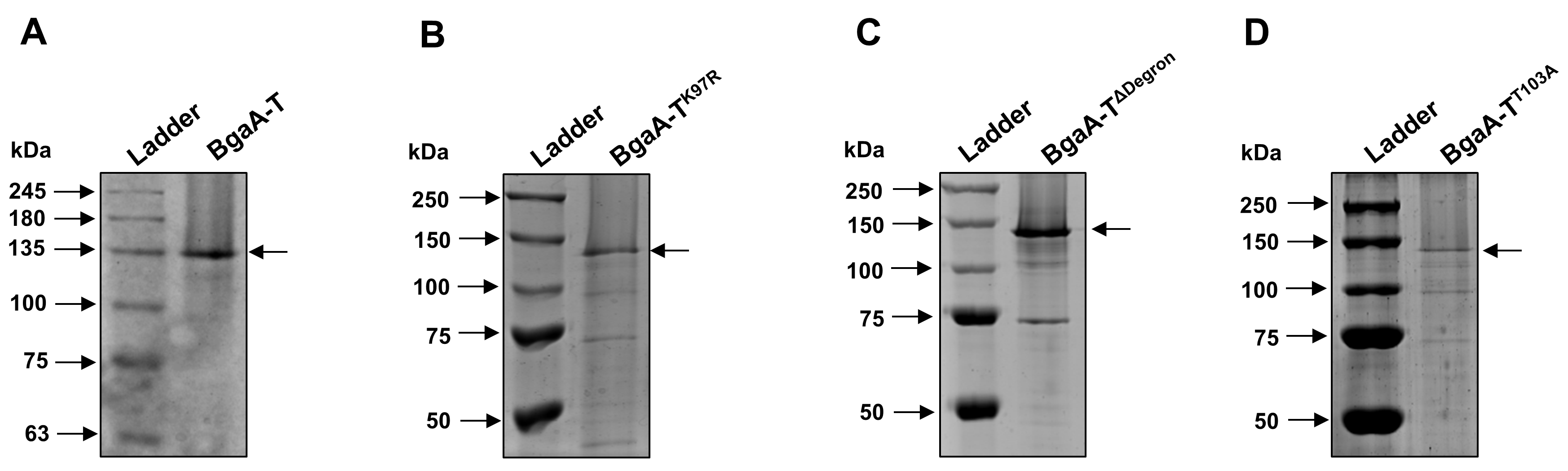

### Fig. S4

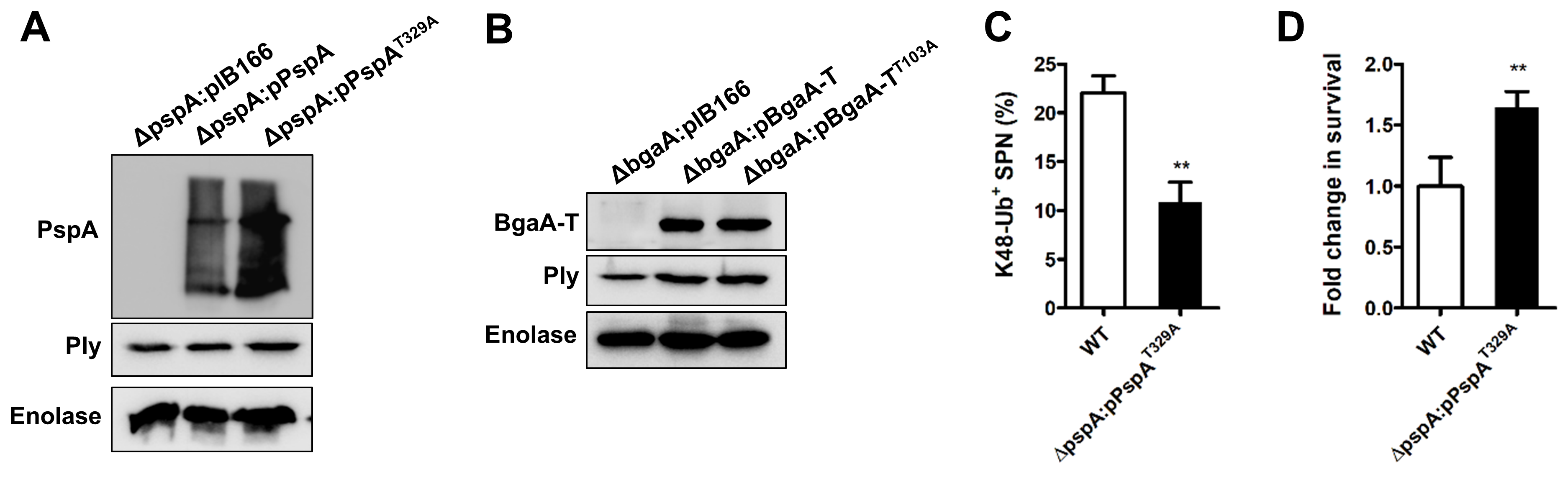
