## Supplementary material for "A novel innate pathogen sensing strategy involving ubiquitination of bacterial surface proteins": Table S1

**Table S1. List of bacterial strains.**

| **STRAIN** | **SOURCE** |
| --- | --- |
| *Streptococcus pneumoniae* R6 (Serotype 2) | Prof. TJ Mitchell, Univ. of Birmingham, UK |
| SPN Δ*bgaA* | This study |
| Δ*bgaA*:pBgaA-T | This study |
| Δ*bgaA*:pBgaA-T^K96R^ | This study |
| Δ*bgaA*:pBgaA-T^K97R^ | This study |
| Δ*bgaA*:pBgaA-T^K96R,K97R^ | This study |
| Δ*bgaA*:pBgaA-T^ΔDegron^ | This study |
| Δ*bgaA*:pBgaA-T^T103A^ | This study |
| Δ*bgaA*:pHysA | This study |
| Δ*bgaA*:pHysA^Degron^ | This study |
| SPN Δ*pspA* | This study |
| Δ*pspA*:pPspA | This study |
| Δ*pspA*:pPspA^K314R^ | This study |
| Δ*pspA*:pPspA^K315R^ | This study |
| Δ*pspA*:pPspA^K314R,K315R^ | This study |
| Δ*pspA*:pPspA^ΔDegron^ | This study |
| Δ*pspA*:pPspA^T329A^ | This study |
| Δ*pspA*:pHysA^Degron^ | This study |
| *Salmonella* Typhimurium (STm) ATCC 14028 | ATCC |
| STmΔ*rlpA* | This study |
