## Supplementary material for "A novel innate pathogen sensing strategy involving ubiquitination of bacterial surface proteins": Table S2

**Table S2. List of primers.**

| **NAME** | **TYPE** | **PRIMER SEQUENCE** |
| --- | --- | --- |
| BgaA-Upstream | F | 5′-TCATATTCTAGAGGTGTAGGTGCCTTCCCAGA-3′ |
|  | R | 5′-TAAAGTGGATCCAAAACCCTCCTTATATTATATTTAGTG-3′ |
| BgaA-Downstream | F | 5′-ACGGTGGGATCCAAATTTTGATACCTTCTTTATCATT-3′ |
|  | R | 5′-TAAAGTCTCGAGCGCTGATGAACCTGAATCAGTC-3′ |
| BgaA-Flanking | F | 5′-GGGGATCTAAATTCTTCATCGGTT-3′ |
|  | R | 5′AACCGATGAAGAATTTAGATCCCC3′ |
| BgaA-T | F | 5′-GCATACGTCAGGGATCCATGGGAAAGGCCATTGGAATCGG-3′ |
|  | R | 5′-ATCAATCTCTAGATTAGTGATGGTGATGGTGATGACCTGTATTTGGTAAA  GGCTTGCTCACATCTACTAA-3′ |
| BgaA-T^K96R^ | F | 5′-GCATCTGAGAGGAAAGAAGATGAAGCCGTAACTCCAAAAG-3′ |
|  | R | 5′-CTTCATCTTCTTTCCTCTCAGATGCAATAGCCTCAGTTG-3′ |
| BgaA-T^K97R^ | F | 5′-TTGCATCTGAGAAGAGAGAAGATGAAGCCGTAACTCCAAAAG-3′ |
|  | R | 5′-GCTTCATCTTCTCTCTTCTCAGATGCAATAGCCTCAGTTG-3′ |
| BgaA-T^K96,97R^ | F | 5′-CTATTGCATCTGAGAGGAGAGAAGATGAAGCCGTAACTCC-3′ |
|  | R | 5′-TTCATCTTCTCTCCTCTCAGATGCAATAGCCTCAGTTGAAC-3′ |
| BgaA-T^Δ102-109^ | F | 5′-AAAGTGTCTGCTAAACCGGAAG-3′ |
|  | R | 5′-GGCTTCATCTTCTTTCTTCTCAGA-3′ |
| BgaA-T^T103A^ | F | 5′-CGTAGCTCCAAAAGAGGAAAAAGTGTCTGCT-3′ |
|  | R | 5′-ACTTTTTCCTCTTTTGGAGCTACGGCTTCATCTTCTTT-3′ |
| HysA^Degron-BgaA^ | F | 5′-GTAACTCCAAAAGAGGAAGCTCTAGGTGGAAACTTAGTTGAT-3′ |
|  | R | 5′-GGCTTCATCTTCCTTGAATGGGTTATCAGTCGTCTTTCG-3′ |
| PspA-Upstream | F | 5′-CGTTAGATATCACAAGTTGTTGCATCG-3′ |
|  | R | 5′-GGAGCCCATATAGTCATTTTCAG-3′ |
| PspA-Downstream | F | 5′-ATTAGGATCCGCCGATTAAATTAAAGCATG-3′ |
|  | R | 5′-TTTTAGAGCTCGATTGAAGGTCGCTTGA-3′ |
| PspA | F | 5′-TCTCGGCTGCCGCTACGGATCCATGAATAAGAAAAAAATGATTTTAACA  AGT-3′ |
|  | R | 5′-ATCTCTTCTAGACTAGTGGTGATGGTGATGATGAACCCATTCACCATTGG  CAT-3′ |
| PspA^K314R^ | F | 5′-AAGCTGACCTTAGGAAAGCAGTTAATGAGCCAGAAAAACC-3′ |
|  | R | 5′-CATTAACTGCTTTCCTAAGGTCAGCTTCAGTTTTTTCTAA-3′ |
| PspA^K315R^ | F | 5′-AAGCTGACCTTAAGAGAGCAGTTAATGAGCCAGAAAAACC-3′ |
|  | R | 5′-GCTCATTAACTGCTCTCTTAAGGTCAGCTTCAGTTTTTTC-3′ |
| PspA^K314R,K315R^ | F | 5′-CTGAAGCTGACCTTAGGAGAGCAGTTAATGAGCCAGAAAA-3′ |
|  | R | 5′-CATTAACTGCTCTCCTAAGGTCAGCTTCAGTTTTTTCTAATTC-3′ |
| PspA^ΔDegron^ | F | 5′-GCACCAGCTGAACAACCAAAACCAGCGCCGG-3′ |
|  | R | 5′-AGCTGGAGCTGGTTTTTCTGGCTCATTAACTGCTTTCT-3′ |
| PspA^T329A^ | F | 5′-CTCCAGAAGCTCCAGCCCCAGAAGCACCAGC-3′ |
|  | R | 5′-GGGGCTGGAGCTTCTGGAGCTGGAGCTGGTTTTTCTGGCTCATTAAC-3′ |
| HysA | F | 5′-CTATATGGATCCATGGACTACAAGGACGACGACGATAAGCAAACAAAA  ACAAAGAAGCT-3′ |
|  | R | 5′-ATTTAATTAGCGGCCGCCTAGTGGTGATGGTGATGATGGTTGTTCTTTCC  TCTACG-3′ |
| HysA^Degron-PspA^ | F | 5′-CCAGAAACTCCAGCCCCAGAATTCAAGGCTCTAGGTGGAAACTTAGTTG  ATATGG-3′ |
|  | R | 5′-TTCTGGAGCTGGAGCTGGTTTTTCTGGGTTATCAGTCGTCTTTCGGAAAT  GTTCGGG-3′ |
| HysA-upstream | F | 5′-TCTAGATCTGCTTCCTTACCGTTGAC-3′ |
|  | R | 5′-GGATCCTAGGAACTAAATCTCAAATTA-3′ |
| HysA-downstream | F | 5′-GGATCCTTTGTTCATCATCTAGATGA-3′ |
|  | R | 5′-CTCGAGTTTCGCCTAAACTACTTCTAT-3′ |
| pET28A- BgaA-T | F | 5′-TCGATCGTCTAGATTTGTTTAACTTTAAGAAGGAGATATACATGG  GGAAAGGCCATTGGAATCG-3′ |
|  | R | 5′-TATCATTGCGGCCGCGGCGTAGTCGGGCACGTCGTAG  GGGTAAAACAGTCTTTTCTTGTCCTT-3′ |
| PspA- Flanking | F | 5′-TGTTGCATCGTAGCTAAGGATTTAT-3′ |
|  | R | 5′-CCCATCTATTCGTTTATTCAC-3′ |
| PspA-Seq | F | 5′-TGCAAAACTTGAAGATCAA-3′ |
| HysA-Seq | F | 5′-CAAGTGACCAATCCTTCTTCTCGTTA-3′ |
| Chloramphenicol Resistance Cassette | F | 5′-AGCCTTGGATCCATGCGTGAGAATGTTACAGT-3′ |
|  | R | 5′-ATCCGGATCCTACAGTCGGCATTATCTCATA-3′ |
| Spectinomycin Resistance Cassette | F | 5′-ATCCGGATCCAATCTGATTACCAATTAGAATG-3′ |
|  | R | 5′-CCGCGGATCCCATATATAATCTAGAATAAAATTAAC-3′ |
| pGEX-4T-FBXW7 | F | 5′-TGGATCCCCGGAATTCATGAATCAGGAACTGCTCTCTG-3′ |
|  | R | 5′- GTCGACCCGGGAATTCTCACTTCATGTCCACATCAAAG-3′ |
